# Functional profiling of ESKAPE viromes uncovers resistance-limiting phage-host dynamics

**DOI:** 10.64898/2026.08.16.745060

**Authors:** Pengwei Li, Quan Liu, Chunfang Deng, Jinren Ni

**Author notes:** Corresponding author: Jinren Ni, Postal address: Peking University, Beijing 100871, P. R. China. These authors contributed equally (Pengwei Li, Quan Liu).

## Abstract

ESKAPE pathogens drive clinical antibiotic resistance and intractable infections, severely compromising antimicrobial therapies. Bacteriophages are promising alternatives to antibiotics, yet their diversity, function and ecological impacts in ESKAPE pathogens remain poorly defined, hindering phage therapy translation. Here, we integrated 11,947 high-quality ESKAPE genomes with global metagenomic viral data to construct a comprehensive non-redundant virome of 14,496 ESKAPE-associated viruses, including four unreported viral clades. We found pervasive competition among mobile genetic elements (MGEs) in the ESKAPE mobilome, where nested MGE architectures empower low-mobility antibiotic resistance genes (ARGs) with horizontal transfer ability to fuel resistance dissemination. Unlike ARG-rich MGEs, ESKAPE phages carry minimal ARGs and antagonize plasmids to constrain ARG propagation, confirming their biosafety for therapy. We further revealed distinct phage-host arms races, typically virulent phages enrich anti-defense genes to evade bacterial immunity, and novel viruses hijack host methyltransferases targeted by CRISPR–Cas systems. This study establishes a systematic ESKAPE virome resource, demonstrates phages’ dual roles in targeting resistant pathogens and curbing resistance spread, and provides mechanistic support for phage therapy clinical application.

## Introduction

Antimicrobial resistance (AMR) represents a major global threat to modern medicine^1^. A significant proportion of resistant infections in clinical settings are caused by a subset of pathogens collectively termed the ESKAPE organism: *Enterococcus faecium*, *Staphylococcus aureus*, *Klebsiella pneumoniae*, *Acinetobacter baumannii*, *Pseudomonas aeruginosa* and other members of the Enterobacterales group^2–4^. These pathogens are frequently implicated in severe nosocomial infections due to multidrug resistance and invasive phenotypes^3^. The World Health Organization has classified ESKAPE pathogens as a priority concern, given their unique resistance profiles and their central role in the global AMR crisis^5^. They share common traits, including adaptation to healthcare environments, versatile resistance mechanisms, and global dissemination of high-risk clones^6^. Their clinical intractability is further compounded by robust biofilm formation, secretion of diverse toxins and enzymes, rapid environmental adaptation, and horizontal gene transfer via conjugation and mobile genetic elements (MGEs), which collectively impede therapeutic intervention^7,8^. Given stagnation in the development of novel antimicrobials, alternative therapeutic strategies are urgently required^6,9–11^.

Bacteriophages (phages), viruses that specifically infect bacteria, have renewed interest as potential therapeutics against pathogens^12,13^. Phages offer high host specificity, minimizing collateral damage to commensal microbiota, low therapeutic dosages, rapid intracellular replication, and potential co-evolution with target bacteria^14–17^. Despite their use since the early twentieth century, phage development was historically overshadowed by antibiotics, particularly outside Eastern Europe^18–21^. The global rise in AMR has reinvigorated interest in phage therapy^21^, though its efficacy and safety depend on comprehensive genomic characterization^7,22^. Challenges include narrow host ranges, emergence of phage-resistant bacteria, phage stability, and effective delivery to infection sites^15,23^. Phage cocktails, combining viruses from diverse families with broad host ranges, may mitigate resistance development^24^, yet this approach requires extensive knowledge of the viral repertoire of the target pathogens^25,26^. Host anti-phage defense systems further influence phage infection dynamics, necessitating consideration of phage–host interactions, including viral countermeasures^27,28^.

To address these knowledge gaps within ESKAPE pathogens and their virome, we performed a large-scale pan-genome analysis of available complete ESKAPE genomes, systematically characterizing their resistome (antibiotic-resistance genes), defensome (anti-phage defense systems), mobilome (MGEs), and virome (bacteriophages). The results reveal the complex interplay between MGEs, resistome, and phage–host interactions, providing critical insights into how mobile elements and viruses shape antimicrobial resistance and phage–host conflicts in ESKAPE pathogens.

## Results and discussion

### Genomic, resistome and defensome landscapes of ESKAPE pathogens

To comprehensively characterize ESKAPE pathogens and their associated virome, we retrieved 11,947 complete genomes from the NCBI RefSeq database, which were dereplicated using an average nucleotide identity (ANI) threshold of 99.9% and a minimum alignment coverage of 30%, yielding 4,538 non-redundant genomes with broad global representation (Fig. 1A, Supplementary Table 1). Comparative genomic analyses revealed substantial variation among ESKAPE species (Fig. 1B). Gram-positive *E. faecium* and *S. aureus* exhibited smaller genome sizes, lower GC content, and reduced coding density relative to Gram-negative *K. pneumoniae, A. baumannii*, *P. aeruginosa*, and *Enterobacter* spp. Notably, *K. pneumoniae* and *Enterobacter* spp., both Enterobacteriaceae members, displayed highly similar genomic features. These genomic differences correspond closely to species-specific functional profiles and virome composition.

**Fig. 1.**
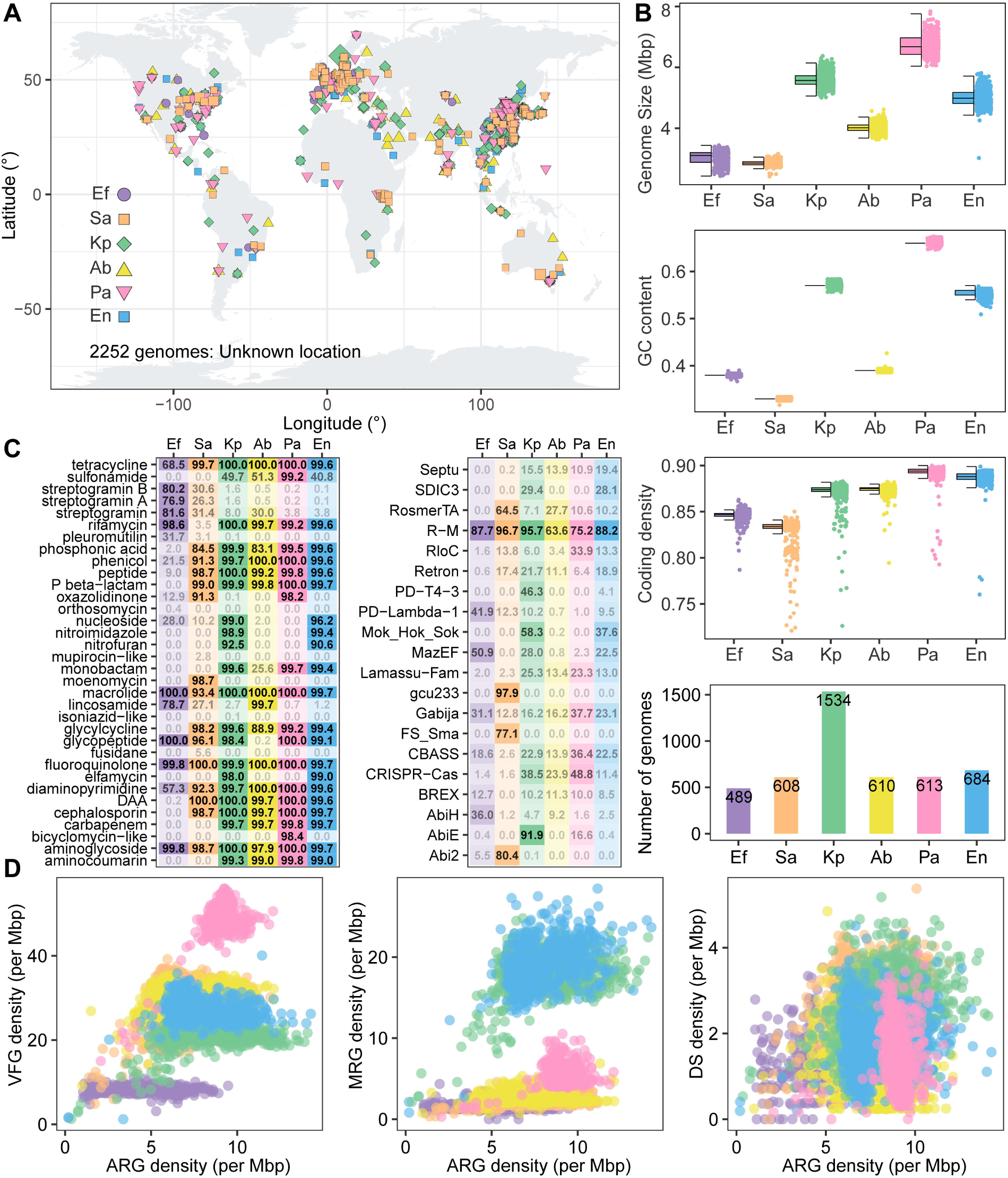
Resistome and defensome landscape of ESKAPE pathogens. **(A)** The geographical location of collected ESKAPE complete genomes. 2,252 genomes with unknown location were not denoted in the map. **(B)** Genomic characteristics (genome size, GC content, and coding density) of all ESKAPE complete genomes. **(C)** The ARG (typed as drug class) and defense system (DS) prevalence within ESKAPE lineages. The value means the proportion of genomes carrying corresponding ARG or DS. The color saturation is correlated with the proportion. In DS heatmap, only the major DS types (In ESKAPE, the sum of prevalence rates exceeds 50) were shown for easy visualization. **(D)** The distribution and relationship of ARG, VFG, MRG, DS (normalized to genome size) in ESKAPE pathogens, different lineages are denoted with different colors. Ab, *A. baumannii*; Ef, *E. faecium*; En, *Enterobacter* spp.; Kp, *K. pneumoniae*; Pa, *P. aeruginosa*; Sa, *S. aureus*.

To investigate antimicrobial resistance (resistome) and anti-phage defense mechanisms (defensome), we annotated antibiotic resistance genes (ARGs), defense systems (DSs), virulence factor genes (VFGs), and metal resistance genes (MRGs) (Supplementary Table 2, 3). ARGs were grouped by antibiotic class, and prevalence was calculated as the proportion of genomes harboring each gene (Fig. 1C). ARG distribution was heterogeneous across species. Resistance to orthosomycin, mupirocin-like, isoniazid-like, and fusidane antibiotics was rare, suggesting retained efficacy against ESKAPE pathogens, whereas tetracycline, macrolide, fluoroquinolone, and aminoglycoside resistance was widespread, indicating limited clinical utility. Certain ARGs exhibited patterns specific to Gram type: sulfonamide, carbapenem, and aminocoumarin resistance were absent in Gram-positive species but prevalent in Gram-negative pathogens, whereas streptogramin resistance was largely restricted to Gram-positive species. These patterns highlight potential avenues for narrow-spectrum therapeutics^11^. The distribution of DSs was similarly heterogeneous, with restriction–modification (R–M) systems predominating across all species (Fig. 1C). Unlike ARGs, DSs displayed no consistent species-specific pattern, indicating that anti-phage defense repertoires are more dynamic and flexible. Even closely related species, such as *K. pneumoniae* and *Enterobacter* spp., show comparable resistome but less consistent DS profiles.

Both ARGs and DSs are adaptive traits that mitigate environmental threats—antibiotics and phages—but impose metabolic costs, potentially creating trade-offs^29,30^. Both gene types are frequently mobilized via MGEs^31,32^, and MGE cargo limitations may bias gene carriage^33^. Certain DSs can also restrict MGE mobility, indirectly limiting ARG dissemination^34^; analogous dynamics may affect MRGs and VFGs^8^. To explore the relationships among these functional traits in ESKAPE pathogens, we performed stringent functional annotation (see Methods) and analyzed their species-specific carriage patterns (Fig. 1D). *P. aeruginosa* encoded higher abundances of ARGs, MRGs, and VFGs compared with *E. faecium* and *S. aureus*. Correlation analyses of ARG–VFG and ARG–MRG abundances clustered species distinctly, whereas DS–ARG relationships were scattered, showing no clear correlation. These findings further underscore the flexible and variable nature of DSs, emphasizing that interpretations of their interplay with other adaptive traits require consideration of ecological and evolutionary context^35^.

### MGE competition and nested mobility shape ARG dissemination in ESKAPE pathogens

Bacteria rapidly adapt to fluctuating environments through horizontal gene transfer (HGT)^36^, facilitated by mobile genetic elements (MGEs), which are broadly classified into plasmids, bacteriophages (phages), and integrative elements^37^. Plasmids are typically extrachromosomal^8^, phages may exist as free particles or integrated temperate forms (prophages), and integrative elements are generally stably embedded in the chromosome. Autonomous integrative elements can excise and reintegrate at alternative loci via transposases, as in insertion sequences (ISs) and transposons (Tn)^38^, or via integrase/excisionase systems, as in integrative and conjugative elements (ICEs) and integrative and mobilizable elements (IMEs)^39^.

Using MGE-specific detection pipelines, we identified 35,312 MGEs in ESKAPE genomes: 10,199 plasmids, 18,411 prophages, 4,799 ICEs/IMEs, 1,594 integrons, 309 ISs, and 84 composite transposons (Tn) (Supplementary Table 4–8). Plasmids were prevalent in all ESKAPE species except *P. aeruginosa* and *S. aureus* (Fig. 2A). Interestingly, we observed an inverse relationship between plasmid and prophage prevalence: except in *P. aeruginosa*, increases in plasmid abundance corresponded with decreases in prophage abundance, and vice versa. *E. faecium* genomes exhibited the highest plasmid and ICE/IME prevalence but the lowest prophage abundance, suggesting that competitive interactions between plasmids and prophages may drive this pattern.

**Fig. 2.**
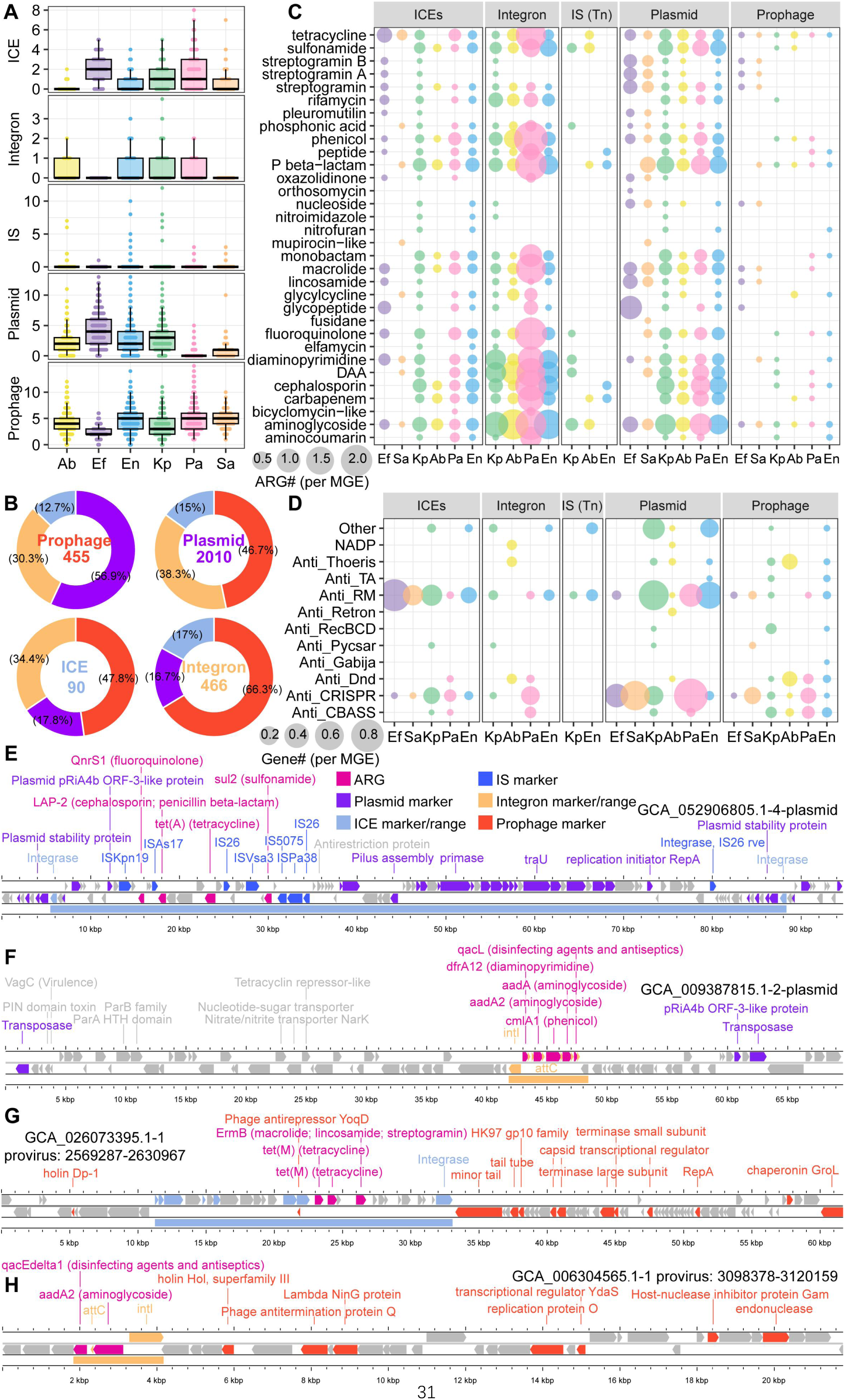
MGE competition and nested parasitism/symbiosis shape the resistome of ESKAPE. **(A)** Number of MGEs in genomes of different ESKAPE lineages. **(B)** The proportion of CRISPR spacer targeted MGE type (prophage, plasmid, ICE, and integron), which reflecting the competition between MGEs. The text colors correspond to the image colors. **(C)** The number (normalized to corresponding MGE number) of ARGs (typed as drug class) in ESKAPE MGEs. **(D)** The number (normalized to corresponding MGE number) of anti-defense genes in ESKAPE MGEs. **(E–H)** Representative MGEs contribute to the horizontal transfer of ARGs through nested parasitism/symbiosis. **(E)** Composite transposons (composed of ISs) that harbor ARGs, which are located in an ICE. The ICE is then carried by a plasmid. **(F)** An integron harboring ARGs is located in a plasmid. **(G)** An ICE harboring ARGs is located in a prophage. **(H)** An integron harboring ARGs is located in a prophage.

CRISPR–Cas systems provide adaptive immunity by capturing fragments of foreign DNA as spacers^40–42^. Aligning spacers to protospacers in other MGEs can reveal historical interactions. Analysis of CRISPR spacers (Supplementary Table 9) across plasmids, prophages, integrons, and ICEs/IMEs showed that 56.9% of prophage-encoded spacers targeted plasmids, whereas roughly half of plasmid, ICE/IME, and integron spacers targeted prophages (Fig. 2B, Supplementary Table 10). These findings indicate substantial reciprocal targeting and support competitive interactions between plasmids and prophages in ESKAPE genomes. We further examined MGE contributions to the resistome. Prophages carried far fewer ARGs than other MGEs, with contributions 10- to 1,000-fold lower than plasmids and integrative elements^43^ (Fig. 2C). In contrast, anti-defense genes and defense systems were encoded at comparable levels across MGEs (Fig. 2D), highlighting that prophages rarely serve as ARG reservoirs.

Not all MGEs are intrinsically mobile across strains: ISs, transposons, integrons, and IMEs are generally restricted to intra-genomic mobility^37^. However, when embedded within conjugative plasmids or prophages, these elements may acquire horizontal transfer potential via plasmid conjugation or phage transduction. We identified examples of nested MGE arrangements that facilitate inter-strain ARG dissemination (Fig. 2E–H). For instance, composite transposon–encoded ARGs were embedded within a putative IME, itself integrated into a plasmid (Fig. 2E); similarly, integron-borne ARGs were found within plasmids or prophages (Fig. 2F, H). Such hierarchical MGE architectures enable the horizontal transfer of otherwise immobile ARGs, extending their dissemination beyond vertical inheritance. These findings underscore the multilayered contributions of MGEs to the ESKAPE resistome, where competition may constrain or promote ARG spread, while nested associations mechanistically facilitate mobilization of non-transferable resistance determinants.

### The extensive ESKAPE virome and their limited ARG carriage

The observations that ESKAPE-associated prophage compete with plasmids, rarely carry ARGs, and mediate ARG transfer via transduction at rates approximately 1,000-fold lower than conjugative elements^44^ underscores the potential of phages as therapeutic agents in the post-antibiotic era. Recent metagenomic studies leveraging CRISPR spacer-based approaches have enabled the identification of previously unisolated viruses infecting diverse microbial taxa^45,46^, including Candidate Phyla Radiation (CPR) bacteria^47^, Asgard archaea^48,49^, and DPANN archaea^50^. By systematically profiling CRISPR–Cas systems across ESKAPE genomes and integrating spacer-guided metagenomic screening (Fig. S1, Supplementary Table 9), we sought to illuminate the largely uncharted ESKAPE virome, revealing both its diversity and its potential implications for antimicrobial strategies.

Consistent with previous reports, ESKAPE pathogens harbor relatively limited CRISPR–Cas systems^3,51,52^ with distinct subtype distributions. Type I-C, II-A, III-A, and IV-A3 systems were exclusively detected in *P. aeruginosa*, *E. faecium*, *S. aureus*, and *K. pneumoniae*, respectively, whereas type I-E systems were identified in *K. pneumoniae* and *Enterobacter* spp., both members of the Enterobacteriaceae family. Notably, type IV CRISPR–Cas systems were consistently associated with conjugative MGEs, particularly plasmids (Fig. S2)^40^. The IV-A3 subtype, previously proposed to function as a plasmid-encoded mediator of inter-MGE competition^40,53^, was detected exclusively in *K. pneumoniae* plasmids in this study (Fig. S2B), suggesting a lineage-specific role in regulating inter-MGE conflicts.

To characterize viruses associated with ESKAPE pathogens, we integrated data from the Virus-Host DB, IMG/VR v4 database, VIRE database, and the collection of ESKAPE genomes (see Methods). Isolated phages were primarily retrieved from the Virus-Host DB^54^, which compiles complete viral genomes deposited in NCBI RefSeq and GenBank. Previously unisolated phages were identified via CRISPR spacer matching against widely used metagenomic viral repositories. Viral identification applied stringent criteria (see Methods): spacer-protospacer matches required 100% coverage with at most one mismatch, followed by quality assessment with geNomad and CheckV, and supported by multiple independent lines of host prediction evidence. This workflow yielded 14,496 non-redundant viral operational taxonomic units (vOTUs), including 9,145 high-quality or complete genomes (Fig. 3A, Fig. S5, Supplementary Table 11). Taxonomic classification (Fig. 3B) revealed that the majority (97%) belonged to Caudoviricetes, whereas a minor fraction (2.8%) was assigned to Faserviricetes. Phages infecting different host show distinct genomic and protein traits (Fig. S3), including GC content, average molecular weight, carbon/nitrogen/sulfur atoms per residue side chain (C/N/S-ARSC), and amino acid usage (AAU) patterns, reflecting the adaptation to host environments. Notably, phages infecting *P. aeruginosa* contained a higher proportion of Faserviricetes (19.4%) than those associated with other ESKAPE species. Fewer viral genomes were recovered for *E. faecium* and *S. aureus* (Fig. 3C). This discrepancy is unlikely due to host genome availability, as comparable bacterial genome numbers were analyzed (Fig. 1B); rather, the low prevalence of CRISPR–Cas systems in these species likely constrained spacer-guided viral discovery (Fig. 1C). This underscores that CRISPR-based detection strongly shapes current ESKAPE virome reconstructions although its higher accuracy, leaving substantial viral diversity uncharacterized, which need further investigation.

**Fig. 3.**
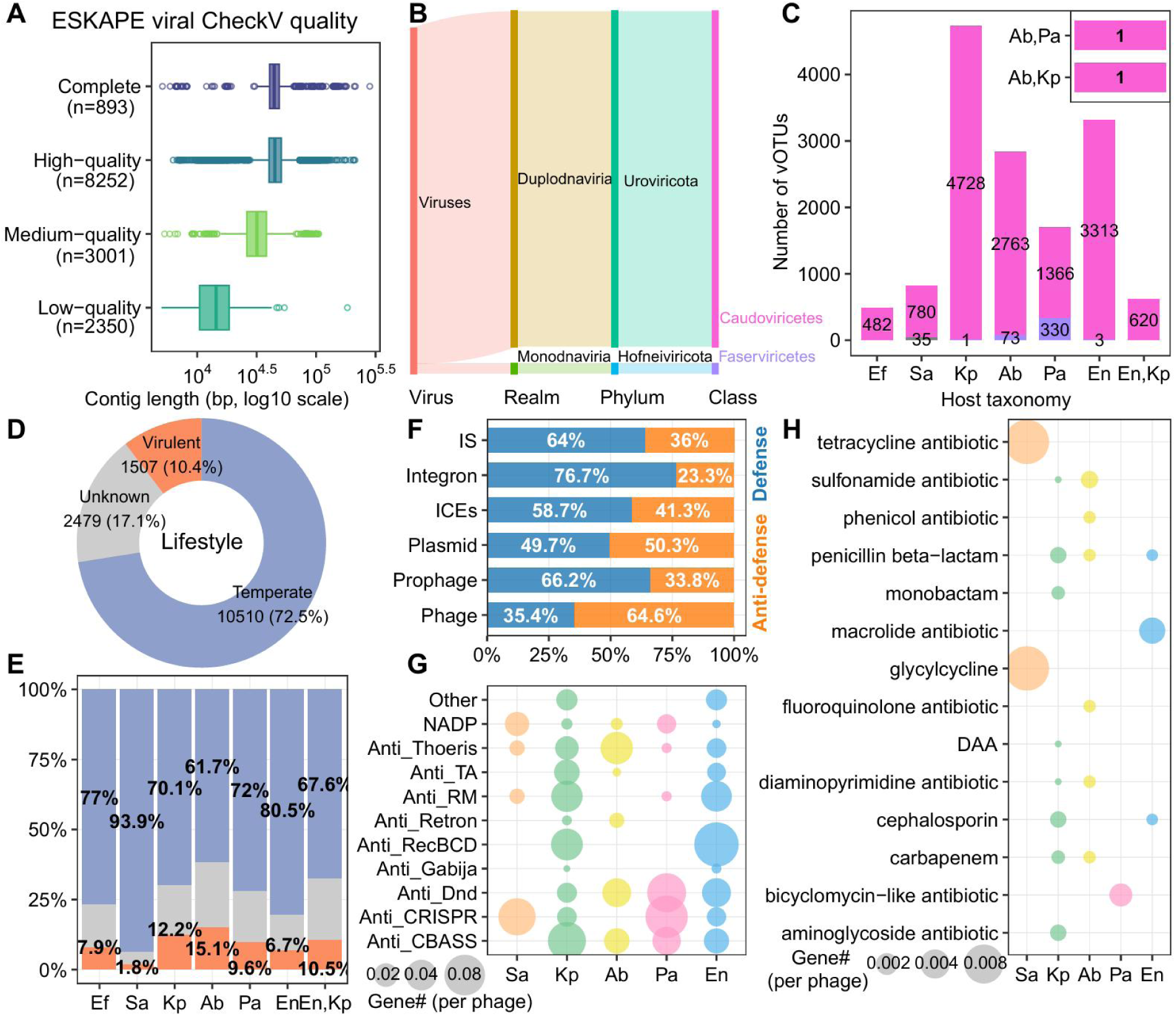
Expansive ESKAPE virome and its limited resistome contribution. **(A)** The CheckV quality of ESKAPE virus. **(B)** Taxonomic annotation of all retained ESKAPE viruses. **(C)** Host details of all retained ESKAPE viruses. **(D)** Predicted lifestyles of all retained ESKAPE viruses. **(E)** The proportion of viral lifestyles. **(F)** The proportion of defense systems vs. anti-defense systems across major MGE, prophage, and phage (don’t include the prophages derived from ESKPAE genomes). **(G)** The number (normalized to corresponding phage number) of anti-defense systems in ESKAPE phages. **(H)** The number (normalized to corresponding phage number) of ARGs (typed as drug class) in ESKAPE phages. DAA: disinfecting agents and antiseptics.

Viruses recovered from metagenomic datasets were primarily derived from human-associated and built environments (Fig. 4A, B), consistent with the ecological distribution of ESKAPE pathogens. Protospacers in these viruses were predominantly associated with nucleotide metabolism, DNA methylation, and viral structural genes (Fig. 4C), reflecting host CRISPR–Cas targeting biases and highlighting viral genes critical for successful infection and lysis^55^. Insights from these genes may guide the engineering of phage components capable of evading ESKAPE CRISPR–Cas defenses.

**Fig. 4.**
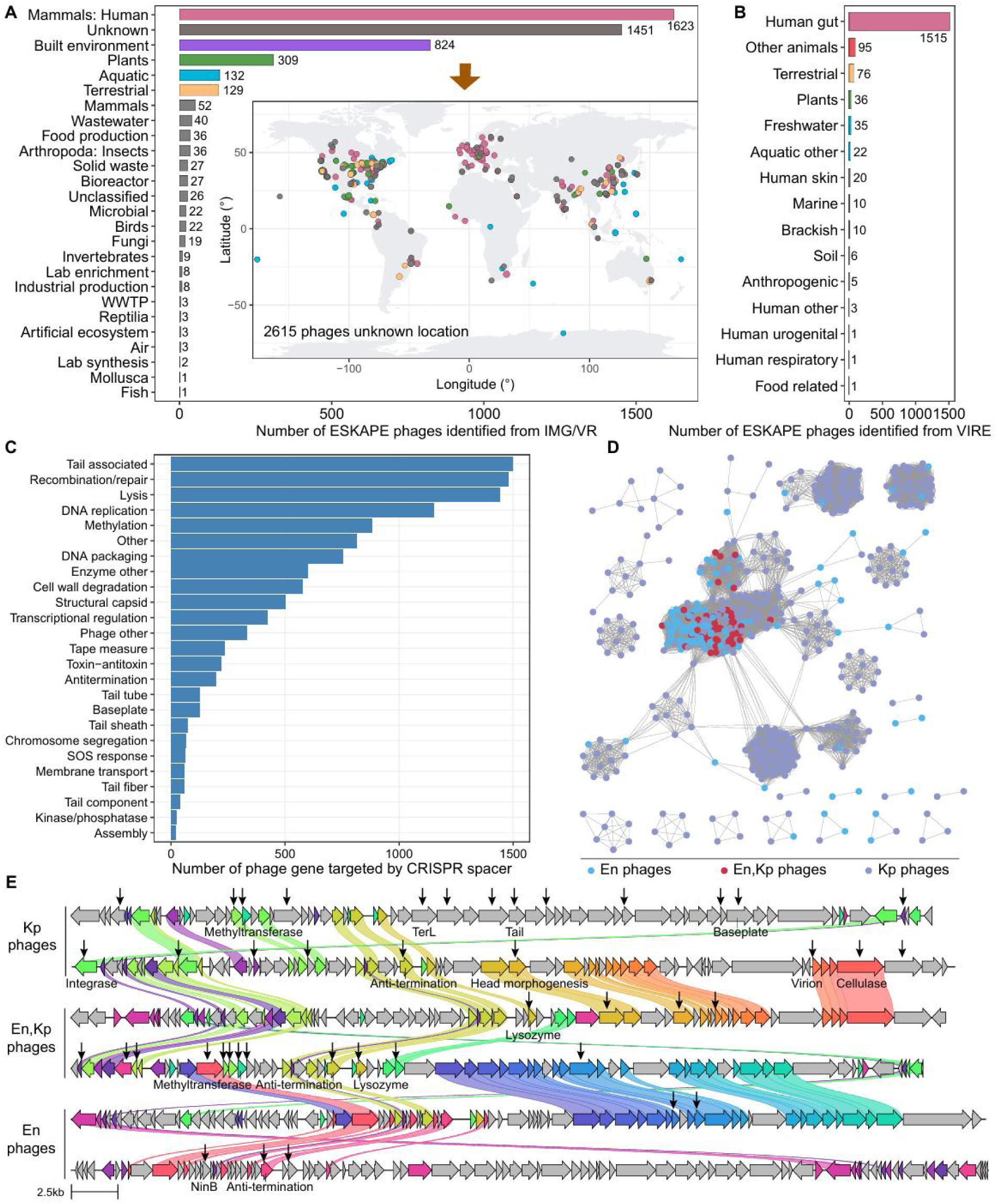
ESKAPE phages exhibit diverse distribution and CRISPR-targeted interactions. **(A)** Ecosystem types that ESKAPE viruses identified from IMG/VR database are sourced from and their geographical distribution. **(B)** Ecosystem types that ESKAPE viruses identified from VIRE database are sourced from. **(C)** The functional types of ESKAPE viral genes targeted by host CRISPR spacers. Viruses identified from IMG/VR and VIRE based on spacer-match methods were included here. **(D)** The protein network of multi-host viruses of Enterobacter spp. and *Klebsiella pneumoniae*, their hosts belong to same family but different genus. **(E)** Comparisons of the genome maps of representative phages of Enterobacter spp. and *Klebsiella pneumoniae*. The genes targeted by host spacers were denoted with arrows. Homologous genes (>30% identity) are highlighted using the same color and linked via shadings.

A subset of 622 vOTUs (4.3%) was predicted to infect multiple host lineages (Fig. 3C). Most (620 vOTUs, including 47 complete genomes) were shared between *K. pneumoniae* and *Enterobacter* spp., consistent with their close phylogenetic relationship and indicative of cross-genus infection. Two additional vOTUs were predicted to infect either *A. baumannii* and *P. aeruginosa* (cross-family) or *A. baumannii* and *K. pneumoniae* (cross-order); however, these genomes were incomplete and were therefore not going deeper analysis. Network analysis (Fig. 4D) showed that multiple-host phages were closely linked with viruses from both hosts, and representative genomic maps revealed high gene homology (Fig. 4E). Interestingly, homologous genes in two phages differed in CRISPR targeting, with some (e.g., cellulase, lysozyme, anti-termination genes) targeted in one but not the other, reflecting the evolutionary plasticity of viral genomes and the challenges hosts face in achieving permanent immunity.

Viral lifestyle prediction indicated that most phages were temperate (72.5%), although proportions varied among ESKAPE species (Fig. 3D, E). Comparative profiling of defense systems and anti-defense genes revealed that virulent phages harbored significantly higher proportions of anti-defense genes than other MGEs, nearly twofold greater than in prophages (Fig. 3F). This likely reflects functional specialization: persistent MGEs and prophages benefit from defense systems to counter competing DNA, whereas virulent phages rely on anti-defense mechanisms to ensure infection, replication, and progeny release^56^. Analysis of DS-to–anti-DS ratios across viral lifestyles further confirmed such functional distinctions (Fig. S4B).

Functional annotation revealed species-specific patterns in anti-defense gene carriage. No anti-defense genes were detected in phages infecting *E. faecium*, and *S. aureus* phages carried fewer anti-defense genes than those targeting other ESKAPE species (Fig. 3G), likely reflecting the limited number of viral genomes recovered. ARGs encoded by ESKAPE phages were similarly rare, occurring at frequencies roughly one order of magnitude lower than anti-defense genes (Fig. 3H), reinforcing the limited role of phages in ARG dissemination within these pathogens.

### Four previously uncharacterized viral lineages expand the ESKAPE virome

All complete ESKAPE viruses identified in this study were compared with previously reported ESKAPE viruses and prokaryotic viruses in the NCBI RefSeq database (release R231) using gene-sharing network analysis (see Methods). Notably, four viral clusters (including singletons represented by a single complete genome; see Methods) exhibited no detectable connections to either RefSeq viruses or previously described ESKAPE viruses (Supplementary Table 12), suggesting that they constitute previously uncharacterized viral taxa within the ESKAPE virome. Comparative genomic analyses revealed conserved gene synteny within each cluster, whereas gene organization differed markedly between clusters (Fig. S6). In proteome-based phylogenetic reconstructions, complete genomes from these four viral groups formed distinct branches clearly separated from reference ESKAPE viruses (Fig. 5A) and from randomly selected other known viruses (blue front in Fig. 5A). These viruses shared an extremely low fraction of orthologous proteins (<10%) and low nucleotide-based intergenomic similarity^57^ with reference viruses and with members of other newly identified clusters (Fig. 5B, Fig. S7).

**Fig. 5.**
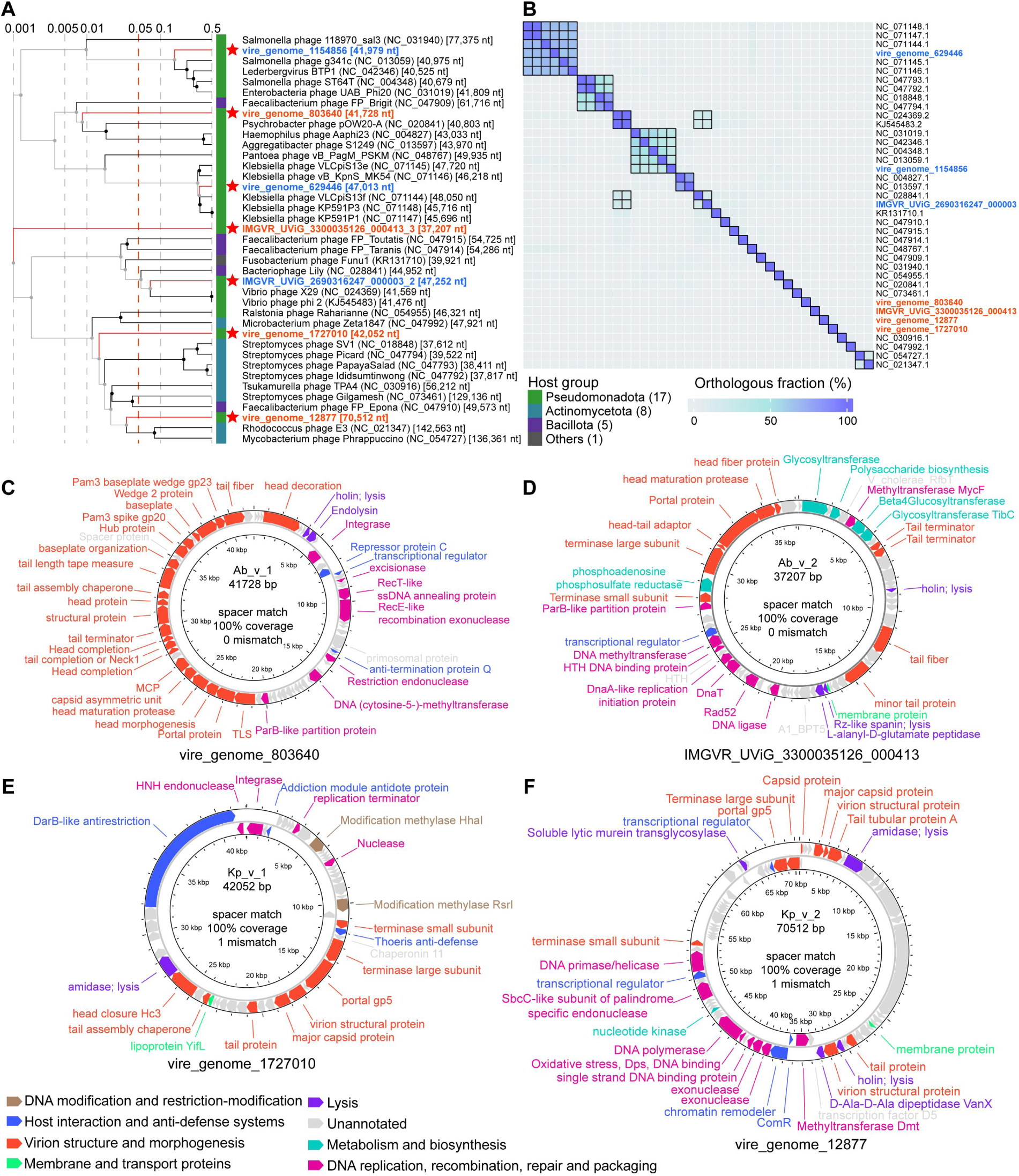
Genome-based taxonomic assignment and complete genomic maps of four new viral groups for *Acinetobacter baumannii* and *Klebsiella pneumoniae*. Whole-proteome-based phylogeny (**A**) and orthologous protein fraction shared by pairwise viruses (**B**) for the taxonomic proposal of novel *Acinetobacter baumannii* and *Klebsiella pneumoniae* viruses. Viruses are indicated by red (vContact3 singletons) or blue (random selected) font. Reference viruses are shown in black font. Reference viruses were selected based on the similarity score of the primary genome-wide proteomic trees constructed using the seven viruses curated in this study and NCBI RefSeq complete viruses. The proteomic trees are based on all-versus-all proteomic similarity matrix and are mid-point rooted. Branch lengths are log-scaled. Red dashed lines in the log-scale phylogenetic trees denote the branch length for family-level demarcation (∼0.05) (**A**). Orthologous protein fractions ≥10% are denoted by the boxes with black borders on the heatmaps (**B**). **(C-F)** The complete circular genomes are evidenced by the presence of DTRs (Supplementary Table 11). Distinct functional categories are differentiated by colors. Proteins not annotated by functional databases are colored grey. For a detailed functional annotation, refer to Supplementary Table 12.

According to established viral demarcation criteria^46,58^, family-level classification in proteome-based phylogenies typically corresponds to branch lengths of ∼0.05, and viruses from different families generally share fewer than 10% of orthologous proteins. From a conservative taxonomic perspective, the current evidence robustly supports the delineation of novel viral clusters or lineages, although formal family-level assignment cannot yet be justified. Accordingly, we provisionally designate these four viral groups as Ab_v_1 (vire_genome_803640), Ab_v_2 (IMGVR_UViG_3300035126_000413), Kp_v_1 (vire_genome_1727010), and Kp_v_2 (vire_genome_12877) (Fig. 5C–F). These groups likely represent the founding members of previously unrecognized viral lineages, potentially corresponding to family-level taxa pending further characterization.

### Novel ESKAPE viruses encode diverse host-interaction functions

Ab_v_1, Ab_v_2, Kp_v_1, and Kp_v_2 belong to the class *Caudoviricetes* within the realm *Duplodnaviria* and encode canonical head-tailed virus proteins, including HK97-fold major capsid proteins (MCPs), terminase large subunits, portal proteins, and tail components (Fig. 5C–F, Supplementary Table 12). These viruses exhibit broad functional potential, particularly in replication, transcription, translation, and host interaction. Many genes could not be annotated using existing databases (see Methods and Supplementary Table 12), highlighting the largely unexplored functional diversity of these novel viruses (Fig. 5).

Ab_v_1 is classified as temperate virus, encoding integrase, which regulates the lysogenic–lytic transition^59^, excisionase, which assists viral DNA excision from the host genome, and a repressor protein that maintains lysogeny by suppressing lytic gene expression.

It may also replicate via recombination- or rolling-circle–based mechanisms, mediated by a 5′–3′ exonuclease for DNA end resection and a recombinase that anneals the resulting 3′-overhang to a homologous strand^60^.

Ab_v_2 lacks canonical temperate markers such as integrase but is likely temperate, encoding a ParB-like partition gene, HTH DNA-binding proteins, and transcriptional regulators, suggesting lysogenic or plasmid-like latent infection with complex regulatory control^61^. It also carries genes involved in metabolism and polysaccharide biosynthesis, typically associated with host cell wall modification during temperate virus–mediated lysogenic conversion, a strategy that prevents superinfection by modifying host receptors^62^.

Kp_v_1 is a temperate virus, encoding integrase and an addiction module antidote protein that, in concert with a toxin, ensures host retention of the phage genome. The addiction mechanism ensures that if the host attempts to discard the phage genome, the host cell perishes due to the rapid metabolism of the detoxification protein coupled with the stability of the toxin. This mechanism constitutes a “subtle hegemony,” compelling the host to maintain a viral lysogenic state^63^. Kp_v_2, in contrast, is likely a lytic virus, lacking integrase and lysogeny maintenance genes, but encoding DNA polymerase, primase/helicase, single-strand DNA-binding proteins, and exonuclease, facilitating rapid hijacking of host replication machinery—a hallmark of virulent phages^64^.

### Viral recruitment of host genes and targeted host immunity reveal phage–host co-evolution

All four viruses have protospacers located within methyltransferase genes, and all were recovered from freshwater metagenomes from distinct geographical locations. Genomic comparison (Fig. 6A) revealed a single conserved gene between Ab_v_1/Ab_v_2 (32% identity) and Kp_v_1/Kp_v_2 (43% identity)—DNA methyltransferase—corresponding to CRISPR-targeted loci. Ab_v_1 and Ab_v_2 were targeted by CRISPR spacers derived from the same *A. baumannii* genome (GCA_052391035.1), isolated from a bovine farm swab sample. Spacer–protospacer alignments exhibited 100% coverage and no mismatches. Kp_v_1 and Kp_v_2 were targeted by spacers from multiple *K. pneumoniae* strains across diverse geographical and ecological contexts, including human blood samples from Houston (USA), Norway, and Germany; a human rectal swab from Australia; and a *Sus scrofa domesticus* cecal sample from Norway. This broad distribution suggests extensive, ongoing interactions, though it may also reflect global dissemination of closely related strains, warranting further epidemiological study.

**Fig. 6.**
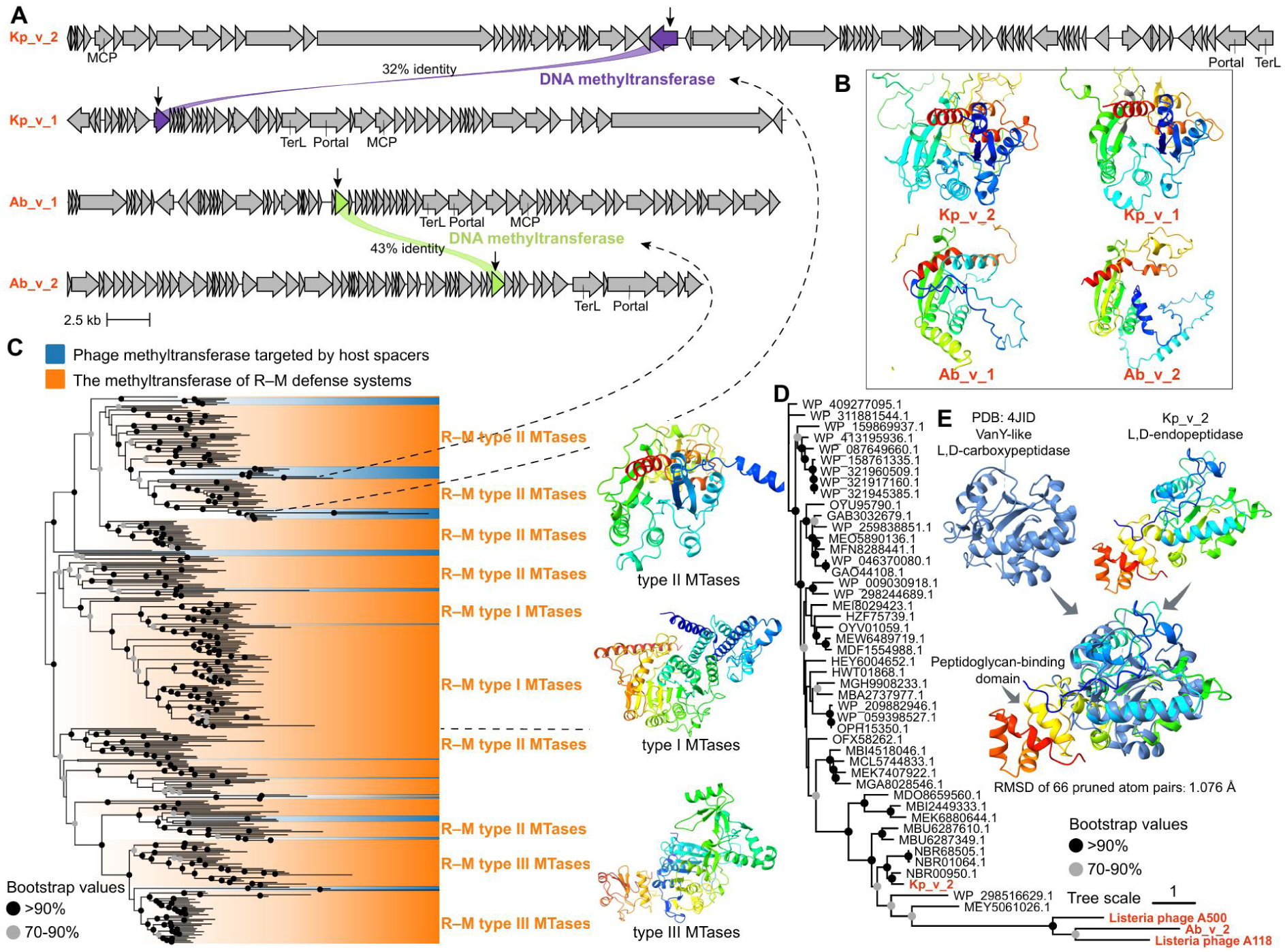
Phylogenetic analysis and structural modelling of genes encoded by the four newly proposed viruses. **(A)** Genomic comparison of four novel ESKAPE viruses. Kp_v_2 and Kp_v_1 are phages of *Klebsiella pneumoniae*. Ab_v_1 and Ab_v_2 are phages of *Acinetobacter baumannii*. The genes targeted by host spacers were denoted with arrows. Homologous genes (>30% identity) are highlighted using the same color and linked via shadings. **(B)** Predicted structure of viral DNA methyltransferase targeted by host CRISPR spacers, corresponding to genes highlighted in panel A. **(C)** Maximum likelihood phylogenies of the DNA methyltransferase, including those encoded by phages and targeted by host spacers and those composed of complete R–M defense systems in ESKAPE genomes. Different R–M types are denoted and the representative protein structures of DNA methyltransferases of type I, type II, type III R–M systems are presented. **(D)** Maximum likelihood phylogenies of the viral L-alanyl-D-glutamate peptidase and homologs (mainly D-alanyl-D-alanine carboxypeptidase) identified from NCBI NR database. **(E)** Comparison of protein structures of VanY-like carboxypeptidase and L-alanyl-D-glutamate endopeptidase identified in Kp_v_2.

Phylogenetic analysis of viral methyltransferases targeted by host spacers and host R–M systems revealed viral acquisition of diverse host MTases (type I–III) (Fig. 6C), indicating viral adaptation to circumvent host R–M defenses, as R–M systems are typically the most dominant defense mechanisms in prokaryotes^31^. Hosts counter this via CRISPR–Cas targeting of viral methyltransferase genes (Fig. 4C, 6A). Additionally, Kp_v_2 and Ab_v_2 encode L-alanyl-D-glutamate peptidases that breaks down the peptidoglycan layer of the bacterial cell wall during the late stages of infection^65^. We find this peptidase is homologous to the zinc-dependent D,D-peptidases VanX, VanY, or VanXY, proteins that play important roles in bacterial resistance to vancomycin^66^. The phylogeny and structural alignment (Fig. 6D, E, Fig. S8) suggest evolutionary recruitment from bacterial proteins to degrade peptidoglycan during late infection. These findings exemplify dynamic virus–host co-evolution, with viruses exploiting host proteins for immune evasion or lysis, countered by host defense mechanisms.

In summary, this study provides a comprehensive, genome-resolved characterization of the resistome, defensome, mobilome, and virome of ESKAPE pathogens. Competitive dynamics among MGEs, compounded by nested parasitism, reveal a complex landscape for horizontal ARG transfer. Viruses rarely carry ARGs, and virulent phages preferentially harbor anti-defense systems, highlighting their therapeutic potential. Viral strategies exploiting host proteins for immune evasion or lysis exemplify the co-evolutionary arms race, counterbalanced by host defenses. Collectively, these findings advance understanding of ARG transmission and phage–host interactions, providing a foundation for optimizing phage therapy against clinically critical ESKAPE pathogens.

## Methods

### ESKAPE pathogens collection

All available complete ESKAPE genomes were retrieved from the NCBI Reference Sequence Database (RefSeq, accessed on 8 December, 2025). Resulting a total of 11,947 complete genomes (921 *Enterococcus faecium*; 2,558 *Staphylococcus aureus*; 4,536 *Klebsiella pneumoniae*; 1,437 *Acinetobacter baumannii*; 1,439 *Pseudomonas aeruginosa*; 1,056 *Enterobacter* spp.). Then these genomes were dereplicated using dRep with ANI ≥ 99.9% and coverage ≥ 30% (parameters: -sa 0.999 -nc 0.30 -comp 50 -con 10). Resulting 4,538 genomes, then the metadata (including available latitude and longitude, and geographical zone) were retrieved from BioSample database. These genomes went through CheckM2^67^ to assess their genome size, GC content, and Coding density.

### Functional annotation

The open reading frames (ORFs) of all MAGs were predicted using prodigal v2.6.3^68^ (-p single) and queried to BacMet2^69^ and virulence factor database (VFDB)^70^ to annotate metal resistance genes (MRGs) and virulence factor genes (VFGs) using diamond v2.1.0^71^ (-k 1 -e 0.00001 --id 60 --subject-cover 80 --query-cover 80). Antibiotic resistance genes (ARGs) were annotated using RGI v6.0.4^72^ against CARD v4.0.1^72^ database, retaining queries annotated as perfect and strict. For antimicrobial peptide (AMP) annotation, we applied same procedure as a previous study^73^, small ORFs were predicted using prodigal v2.6.3^68^ (-p meta -g 11 -n -q) and filtered to retain only complete smORFs (≥33bp ≤303bp partial=00). Then AMPs were predicted using Macrel v1.5.0^74^ (peptides mode). Biosynthetic Gene Clusters (BGCs) were predicted using antiSMASH v8.0.1^75^ (--taxon bacteria --allow-long-headers). Anti-phage defense systems were detected using DefenseFinder v2.0.0^76^. Defense island was defined as arrays of defense genes separated from one another by 10 genes or less and containing at least five genes belonging to a minimum of three different subtypes of defense system, which keep consistent with previous studies^31^.

### Identification of ESKAPE MGEs (plasmid, prophage, integron, ICE/IME and IS)

Plasmid and prophage within ESKAPE genomes were identified and classified using geNomad v1.8.0^77^ (end-to-end). The sequences classified as provirus by geNomad were treated as prophage in later analyses. To lower bias, CheckV was used to trim host regions from the edges of geNomad-identified proviruses to reduce contamination by host genes. Integrons were identified using IntegronFinder v2.0.5^78^ (--local-max --func-annot --gbk --pdf) and complete integrons (Integron with integron integrase nearby *attC* site) were retained for later analyses. Integrative Conjugative Elements (ICEs) and Integrative Mobilizable Elements (IMEs) were detected using ICEfinder v2.0^39^. Insert sequence (IS) was identified using ISfinder^38^ and retained complete ones for later analyses. To identify potential composite transposons, we searched for pairs of IS elements located within 10 kb of each other on the genome. Genes were first functionally annotated across the genome. Those located within predicted MGEs were then identified based on coordinate overlap.

### Validation of CRISPR arrays and identification of CRISPR–Cas systems in ESKAPE genomes

CRISPR arrays in the genomes (including MGEs) were identified using MinCED^79^ and validated with CRISPRCasFinder v4.3.2^80^, retaining arrays with evidence level 4 (repeat conservation index >70% and spacer identity <8%) for subsequent analysis. CRISPR–Cas systems within the contigs containing these CRISPR arrays were annotated with CRISPRCasTyper v1.8.0 (--prodigal meta --no_grid)^81^. Potential MGE competition was analyzed using the spacer matches. The CRISPR spacers extracted from diverse MGEs (plasmids, prophages, integrons, and ICEs/IMEs) were mapped against each other to assess putative inter-MGE targeting using BLASTn^82^ (-evalue 1e-5 -word_size 8 -task blastn-short), retaining matches with 100% coverage and a maximum of one mismatch.

### Identification and collection of ESKAPE viruses

ESKAPE viruses were identified from the IMG/VR v4 database^83^, VIRE database^84^, ESKAPE genomes, and Virus-Host DB^54^ (accessed on 8 December, 2025). Spacers from validated CRISPR arrays were compared to contigs in the IMG/VR v4 and VIRE database^84^ using BLASTn^82^ (-evalue 1e-5 -word_size 8 -task blastn-short), retaining sequences with 100% coverage and a maximum of one mismatch, resulting in 34,565 viral sequences. Provirus sequences within ESKAPE genomes were identified using geNomad v1.8.1^77^ (end-to-end), with further de-contamination through CheckV v1.0.3^85^, resulting 18,427 putative prophage contigs. ESKAPE phage from Virus-Host DB were retrieved by searching the complete species name of ESKAPE, resulting 3,590 viruses. Dereplication of all these viral sequences (56,582) was performed using the CheckV python scripts with 95% ANI and 85% coverage, yielding 17,833 viruses. These viral contigs were further trimmed using geNomad v1.8.0^77^ and CheckV v1.0.1^85^ following previously published criteria^86^. Briefly, for geNomad predictions, contigs of 5-10 kb were required to have a virus score ≥ 0.9, at least one viral hallmark gene, and a virus-marker enrichment > 2.0, whereas contigs ≥ 10 kb were required to have a virus score ≥ 0.8 and at least one hallmark gene or a virus-marker enrichment > 5.0. Other contigs with direct or inverted terminal repeats (DTRs or ITRs) were also maintained. Then all viruses were further trimmed using CheckV, retaining only contigs with viral_gene > 0. This process yielded a total 16,905 ESKAPE viruses. Although CRISPR spacer matches are currently the most reliable and straightforward approach to link metagenomic viral sequences to their likely hosts, there is also caveats associated with this methodology whereby non-infection interactions can also lead to CRISPR-spacer gain events^87^. To further reduce the false positives, we buttress the results with other lines of evidence using iPHoP v1.4.2^88^, which integrates multiple host prediction strategies (gene flux between viruses and the putative hosts, CRISPR spacer matches, and k-mer composition similarity). Based on iPHoP results, only viruses with host prediction belong to ESKAPE and viruses with complete genome were retained, resulting 14,496 viruses. The presence of direct terminal repeats (DTRs) or inverted terminal repeats (ITRs) in the identified viral sequences was determined using geNomad^77^. Sequences containing high-quality DTRs or ITRs were considered as representing circular or linear complete genomes. Complete viral genomes were visualized using Proksee^89^ to provide a comprehensive view of the genomic features. The genomic features, including genomic size, GC content, average molecular weight, carbon/nitrogen/sulfur atoms per residue side chain (C/N/S-ARSC), amino acid usage (AAU) patterns, and proteome-wide isoelectric point (pI) distributions for complete ESKAPE viruses were calculated according previous studies^46,90^.

To collect as many ESKAPE viruses as possible while ensuring high reliability, except widely used IMG/VR database, we also integrated ESKAPE viruses from VIRE^84^ and Virus-Host DB^54^ (accessed on 8 December, 2025). VIRE (1.7 million high- and medium-quality viral genomes) is a planetary-scale resource of viral genomes recovered from over 100,000 publicly available metagenomes spanning host-associated, aquatic, terrestrial, and anthropogenic environments. Virus-Host DB (45,057 complete viral genomes) organizes data about the relationships between viruses and their hosts, represented in the form of pairs of NCBI taxonomy IDs for viruses and their hosts. covers viruses with complete genomes stored in 1) NCBI/RefSeq and 2) GenBank whose accession numbers are listed in EBI Genomes. The host information is collected from RefSeq, GenBank (in free text format), UniProt, ViralZone, and manually curated with additional information obtained by literature surveys.

Predicted proteins of the identified ESKAPE viruses were functionally annotated using HH-suite3^91^. Multiple sequence alignment of each protein was generated using HHblits v3.3.0 with UniRef30_2023_02 database (three MSA generation iterations; e-value, 1e-6). The resulting MSAs were then searched against a variety of publicly available database, including PDB70_May_2025^92^, PfamA 35.0^93^, NCBI CDD v3.19^94^, PHROG v4^95^, SCOPe70 v2.08^96^, and UniProt-SwissProt-viral70_Nov_2021^97^, using HHsearch v3.3.0 (-Z 250 -loc -z 1 -b 1 -B 250 -ssm 2 -sc 1 -seq 1 -norealign -maxres 32000). The anti-defense genes within ESKAPE viruses were annotated using DefenseFinder v2.0.0^76^ (-a).

### Genome-based taxonomic assignment of viruses

A gene-sharing network was built using vConTACT v3.1.6^98^, containing the complete genomes of ESKAPE viruses identified in this study and prokaryotic viruses from the reference database (release 230). Cytoscape v3.10.3^140^ was used for visualizing the network and genome-scale comparisons were performed using the clinker module of CAGECAT v1.0^99^. Proteome-scale phylogeny for bacterial viruses of the class Caudoviricetes and the complete viral genomes identified in this study were generated using the ViPTree server^100^. To simplify illustration, a subtree was manually created by selecting viral sequences identified in this study and their associated viral sequences from the original tree. Typically, viruses were assigned to the same family if they formed a cohesive group within the vConTACT3 network and appeared as a monophyletic group in the ViPTree analysis with a threshold of 0.05^101^. The orthologous fraction of proteins shared by viral genomes was estimated using CompareM (https://github.com/dparks1134/CompareM) (-evalue 1e-5 -identity 30%), and the input sequences were the same as those for ViPTree analysis. The nucleotide-based intergenomic similarities of newly found complete ESKAPE viruses and reference viruses were assessed using VIRIDIC^57^, which implements the traditional algorithm used by the International Committee on Taxonomy of Viruses (ICTV), Bacterial and Archaeal Viruses Subcommittee, to calculate virus intergenomic similarities.

### Protein structural prediction and visualization

To visualize the active sites in multiple sequence alignment, viral L-alanyl-D-glutamate peptidases were aligned with identified homologs in NCBI NR database (BLASTp 30% identity and 60% coverage) using MAFFT v7.525^102^ L-INS-I and the sequence logos was visualized using WebLogo3^103^. All protein structures are predicted using AlphaFold3^104^ webserver except the protein denoted with PDB ID, and the structural model was visualized using ChimeraX^105^.

### Phylogenetic analysis

Viral DNA methyltransferases targeted by host spacers were extracted and aligned with other DNA methyltransferases derived from ESKAPE complete R–M systems. To reduce redundancy, the two protein sets were dereplicated using MMseqs2^106^ easy-cluster (--min-seq-id 0.6 -c 0.8 --cov-mode 1). The resulting proteins were aligned using MAFFT v7.525^102^ L-INS-I and non-informative columns were removed from the alignment using ClipKIT v2.7.0^107^ (-m gappy -g 0.9). Next, the phylogenetic tree was constructed based on the trimmed alignment using IQ-TREE v2 with the parameters: -m MFP+MERGE -B 1000. The resulting phylogeny was visualized using the Chiplot webserver. The same process was applied to generate phylogenetic trees for viral L-alanyl-D-glutamate peptidases. The best fitting models for phylogenetic reconstructions were Q.pfam+F+R8 and WAG+I+G4, respectively.

## Data Availability

All data support this study are available.

## Code Availability

Wrapper scripts supporting all key analyses of this work are available on GitHub (https://github.com/lianmsu/ESKAPE_mobilome_virome).

## Acknowledgements

This work was supported by the National Natural Science Foundation of China (425B2048, 51721006, and 92047303). Supports from the High-performance Computing Platform of Peking University are acknowledged.

## Author Contributions

J.R.N. designed the research. P.W.L. conducted the bioinformatic and statistical analysis with help of C.F.D.. Q.L. conducted the experimental exploration and validation. P.W.L. and Q.L. wrote the manuscript and J.R.N. revised the manuscript. All the authors read and approved the final manuscript.

## Supplementary Figures

**Fig. S1.**
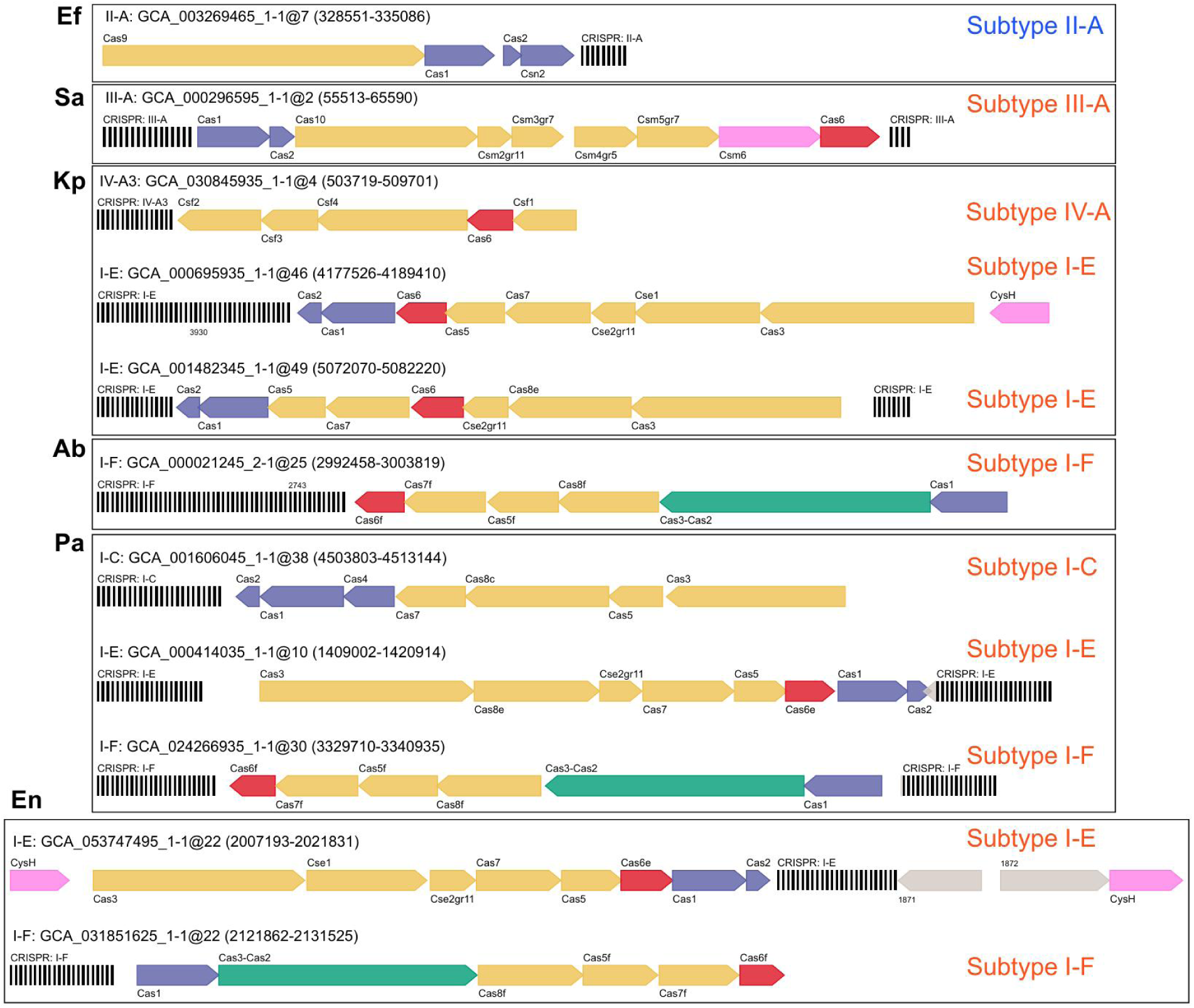
Genetic arrangements of CRISPR–Cas systems in ESKAPE pathogens. Genetic arrangements of each subtype of the identifiable CRISPR–Cas system in ESKAPE pathogens. If one subtype is possessed by multiple genomes for a species, the representative one is shown. CRISPR–Cas system subtypes belonging to classes 1 and 2 are in red and blue text, respectively, on the top right corner within each frame.

**Fig. S2.**
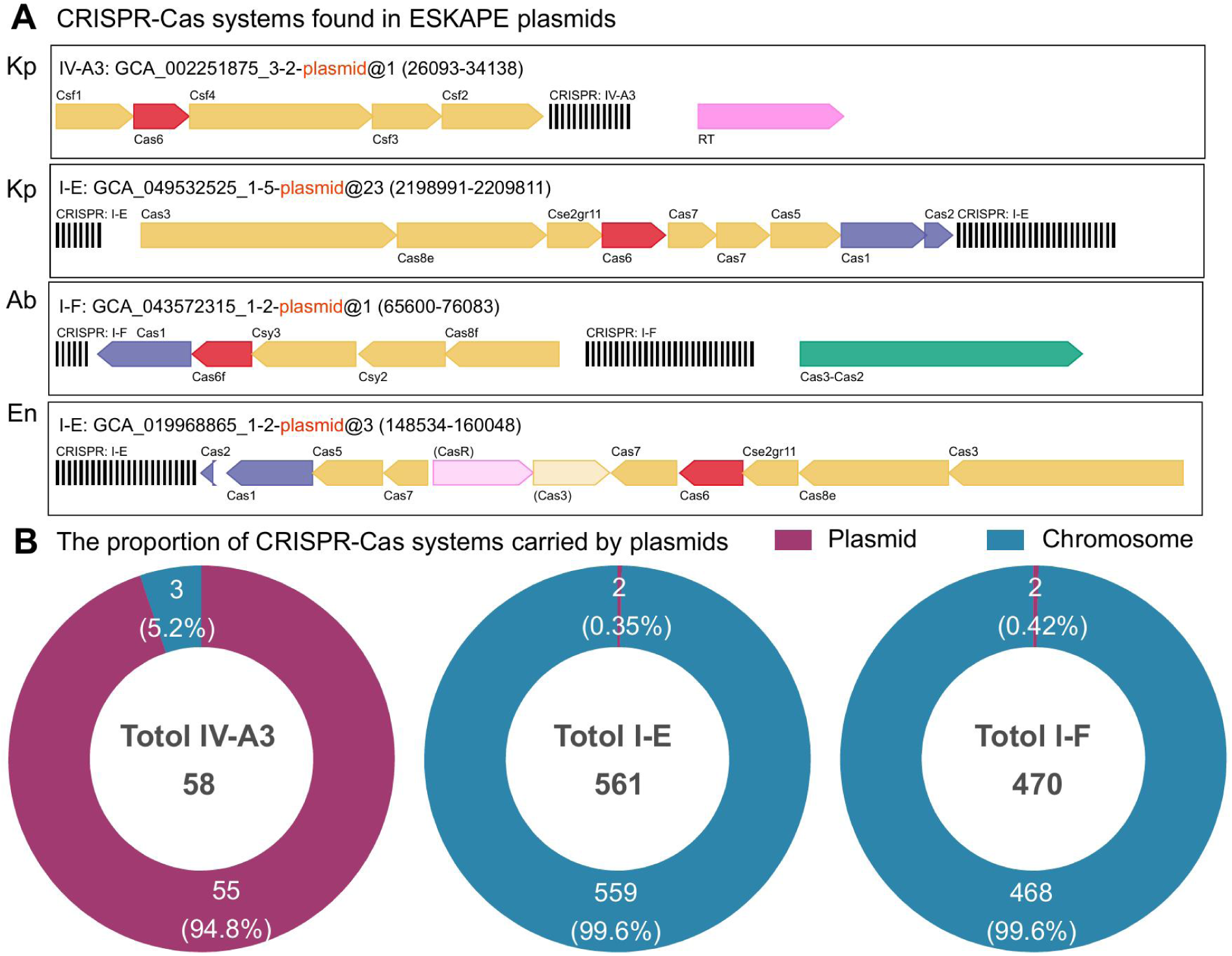
Genetic arrangements of CRISPR–Cas systems in ESKAPE plasmids. **(A)** Genetic arrangements of each subtype of the identifiable CRISPR–Cas system in ESKAPE plasmids. If one subtype is possessed by multiple plasmids for a species, the representative one is shown. **(B)** The proportion comparison of different subtype carried by plasmid or chromosome.

**Fig. S3.**
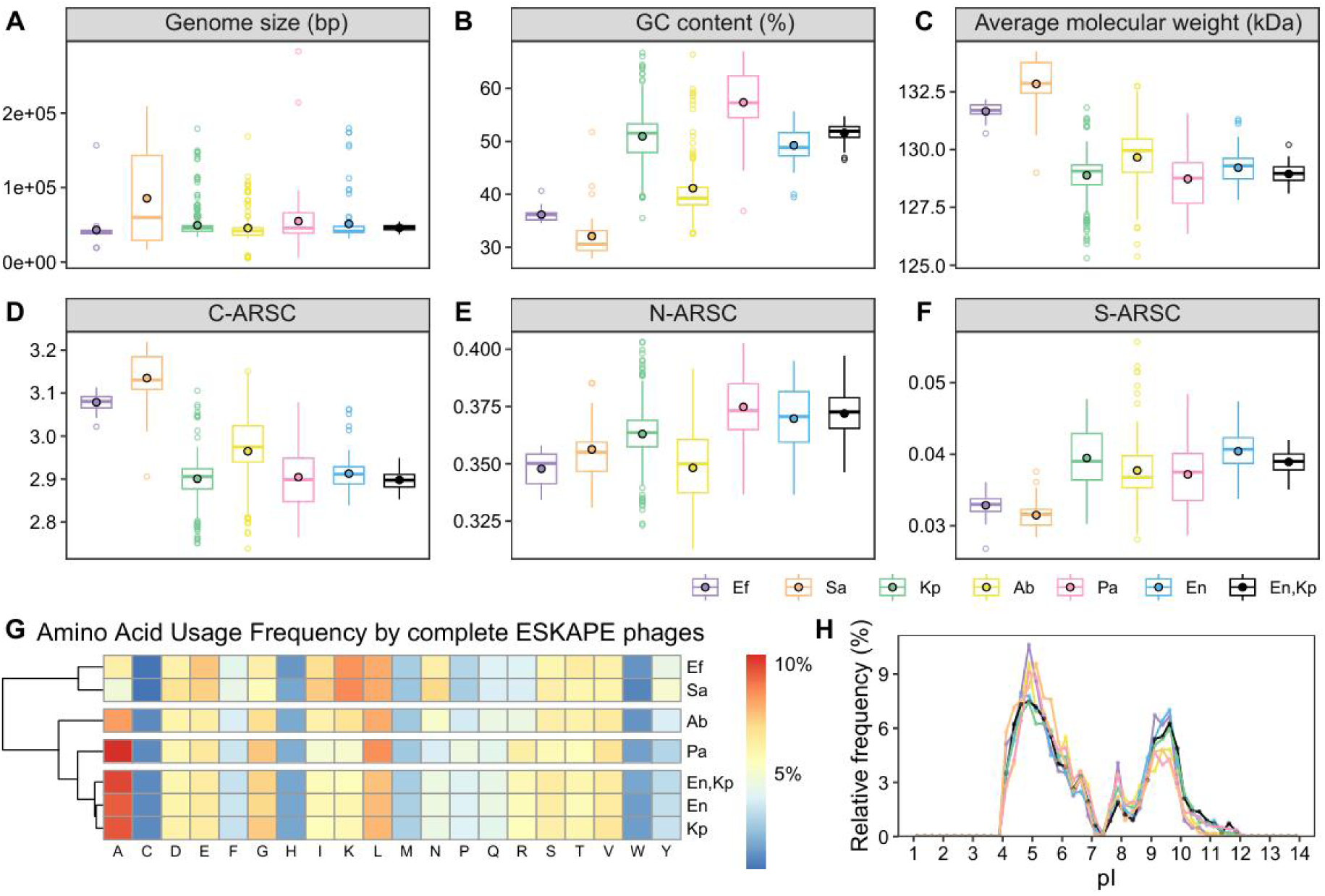
Genomic and protein traits of ESKAPE viruses. Boxplots indicate the difference between complete ESKAPE viruses, specifically with regard to: **(A)** genome size; **(B)** GC content; **(C)** molecular weight; **(D)** C-ARSC; **(E)** N-ARSC; and **(F)** S-ARSC. For each boxplot, central line and whiskers indicate the median and 1.5 times the interquartile range. The upper and lower sides of boxes represent the interquartile range between 25th and 75th percentile. The dot with black border indicates the mean value. Points beyond whiskers are potential outliers. **(G)** Heatmap of amino acid usage patterns. Amino acid frequencies were calculated using proteins encoded by the complete ESKAPE phages. **(H)** Distribution of isoelectric point (pI) values inferred for proteins encoded by the complete ESKAPE phages.

**Fig. S4.**
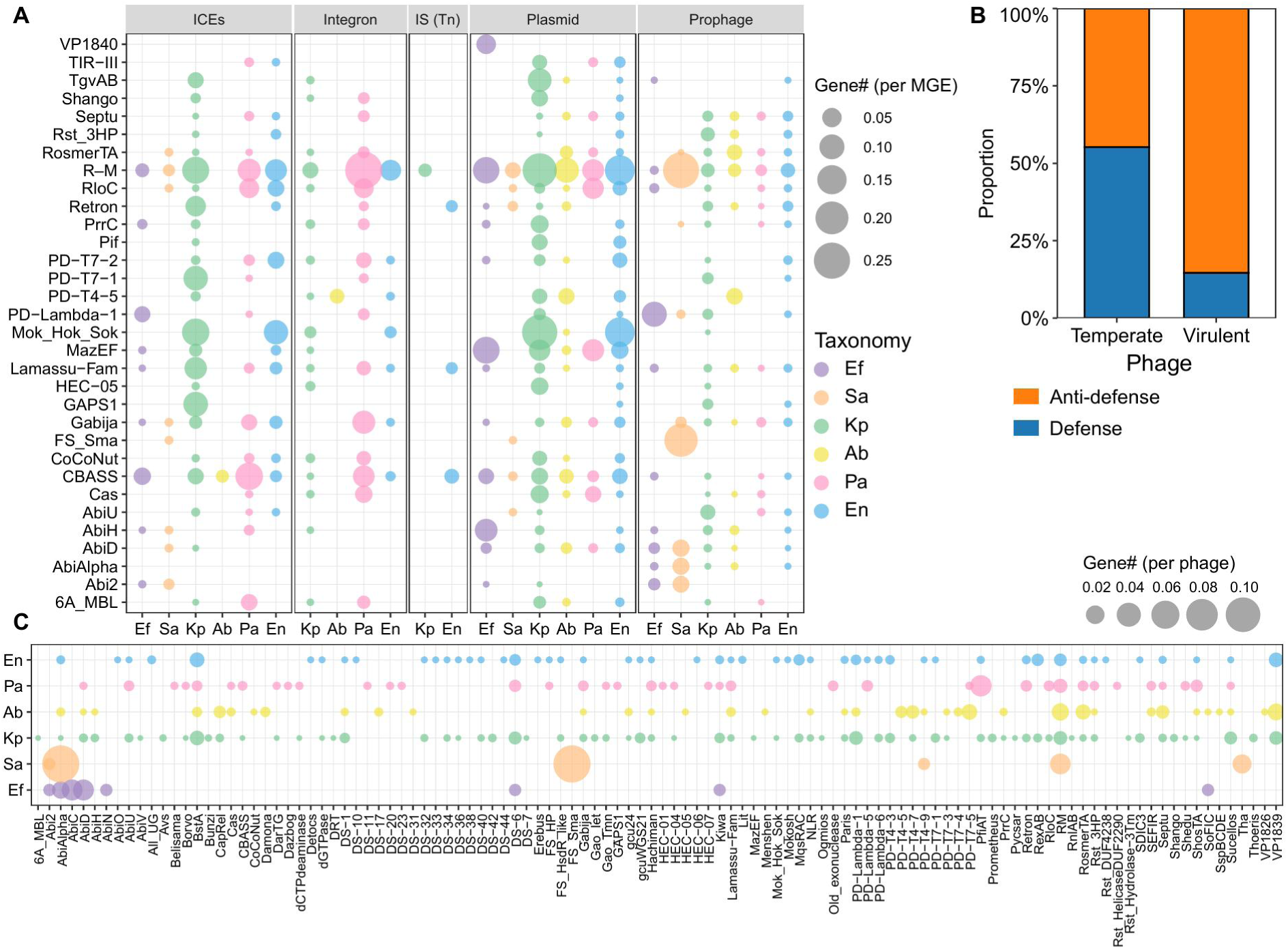
Defense systems in MGEs and ESKAPE viruses. **(A)** The number (normalized to corresponding MGE number) of defense systems possessed by ESKAPE MGEs. For visualization, the primary defense systems are displayed here (with over 110 carried by MGE). **(B)** The proportion of defense systems vs. anti-defense systems across ESKAPE phages when grouped by lifestyles (3091 temperate phages and 313 virulent phages). **(C)** The number (normalized to corresponding phage number, don’t include the prophage identified from ESKAPE genomes) of defense systems in ESKAPE phages.

**Fig. S5.**
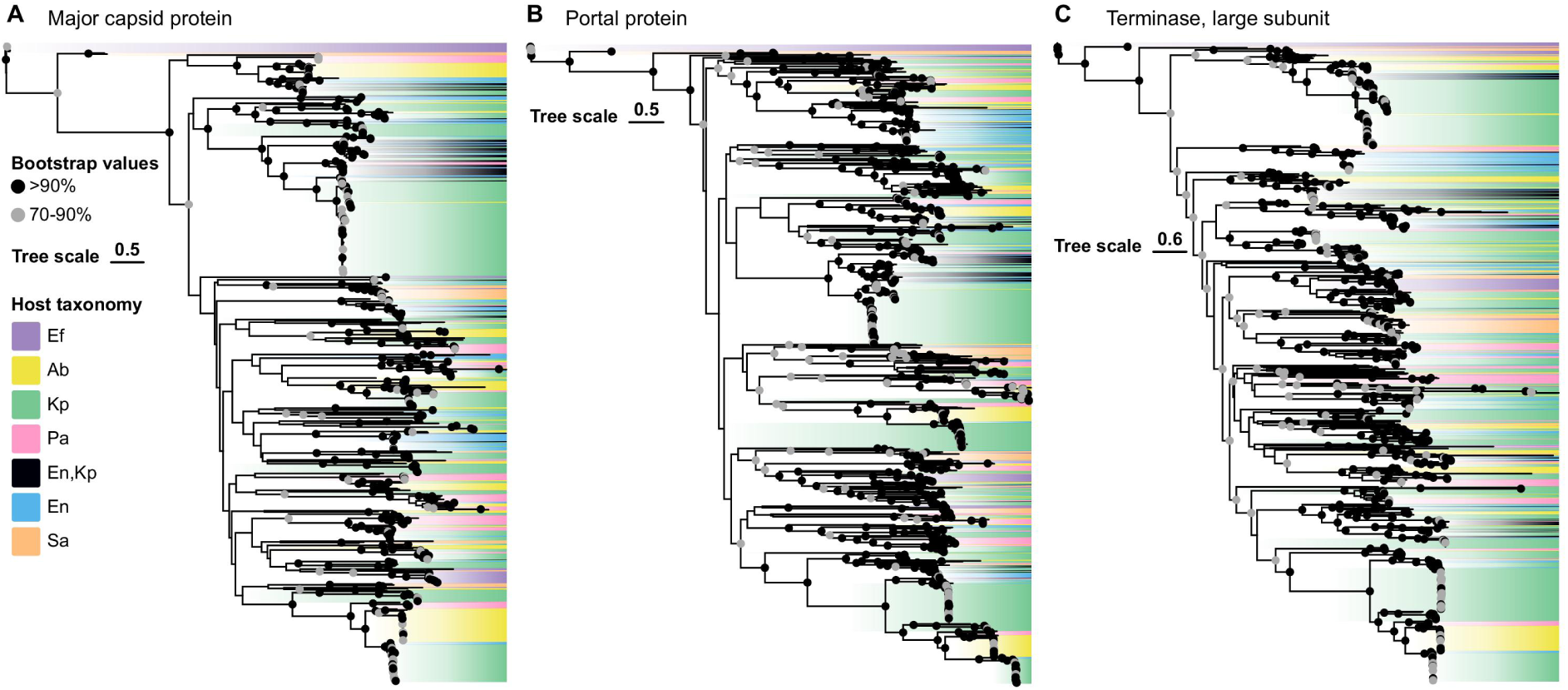
Maximum-likelihood phylogenetic tree of major capsid proteins, portal proteins and terminase large subunits from complete ESKAPE viruses. Bootstrap support values are indicated with nodes. The viral leaves were colored with corresponding host taxonomy.

**Fig. S6.**
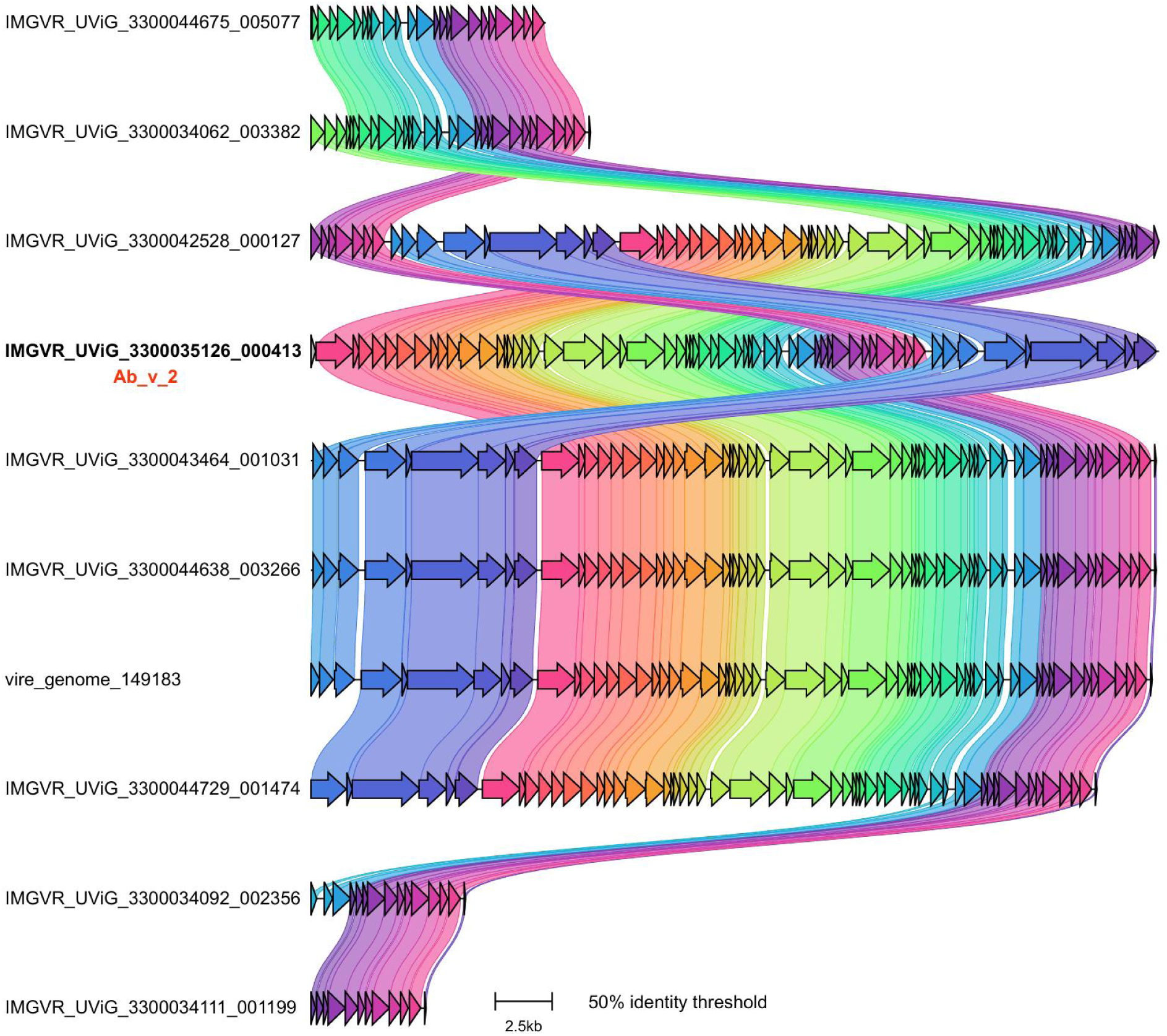
Genome maps showing the close relationships among Ab_v_2 virus group.

**Fig. S7.**
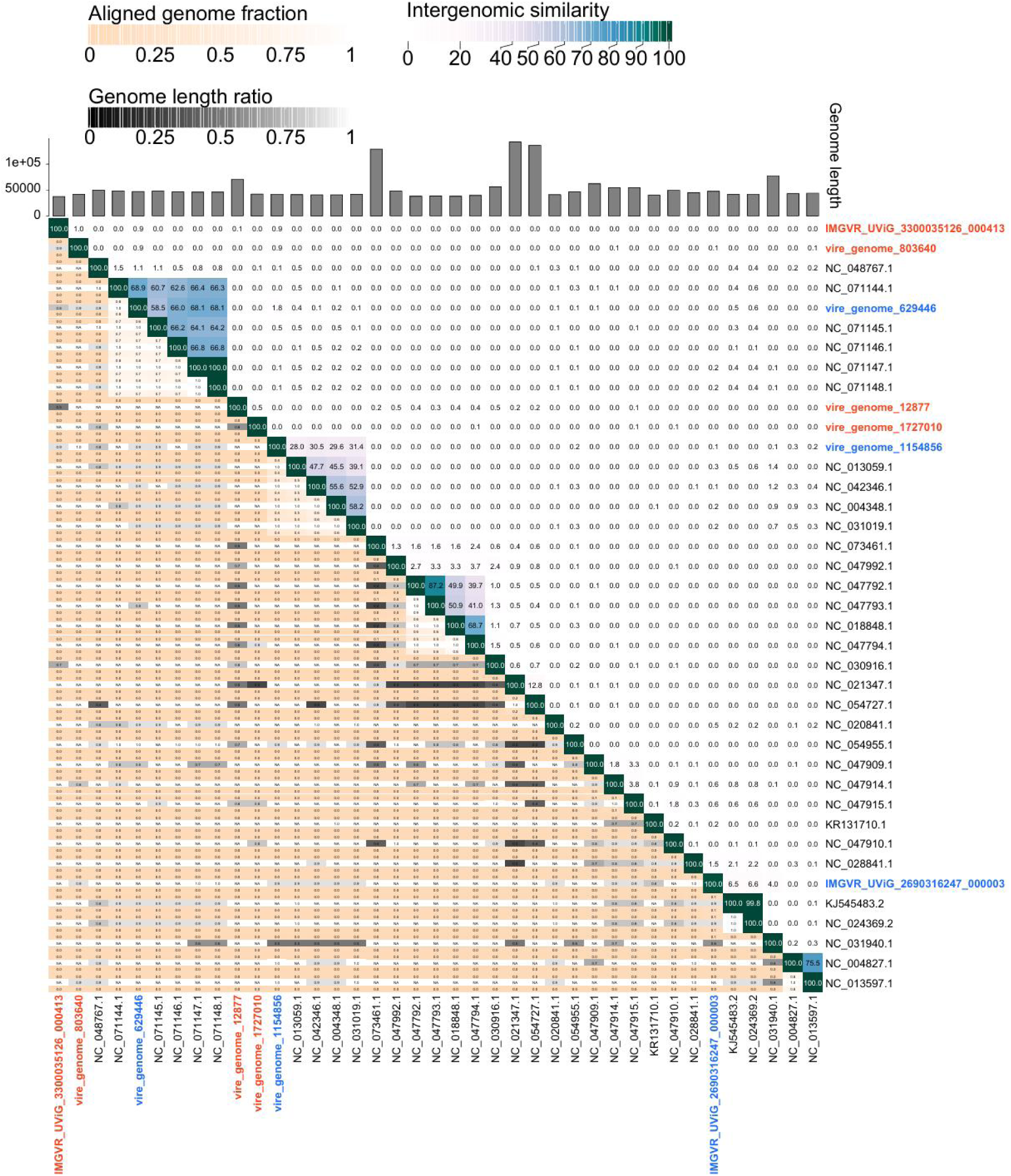
VIRIDIC-generated heatmap depicts intergenomic similarity (right half) and alignment indicators (left half and top annotation). In the right half, a color gradient visualizes phage genome clustering based on similarity—darker colors indicate higher relatedness, with numerical similarity values shown to one decimal place. The left half displays three alignment metrics per genome pair (from top to bottom): the aligned fraction of the row genome, the genome length ratio of the pair, and the aligned fraction of the column genome. Darker shades denote low values — orange-to-white gradients reflect low aligned fractions, while black-to-white gradients highlight substantial length disparities. Aligned fractions inversely correlate with phylogenetic distance; darker colors correspond to low similarity, and lighter colors to high similarity.

**Fig. S8.**
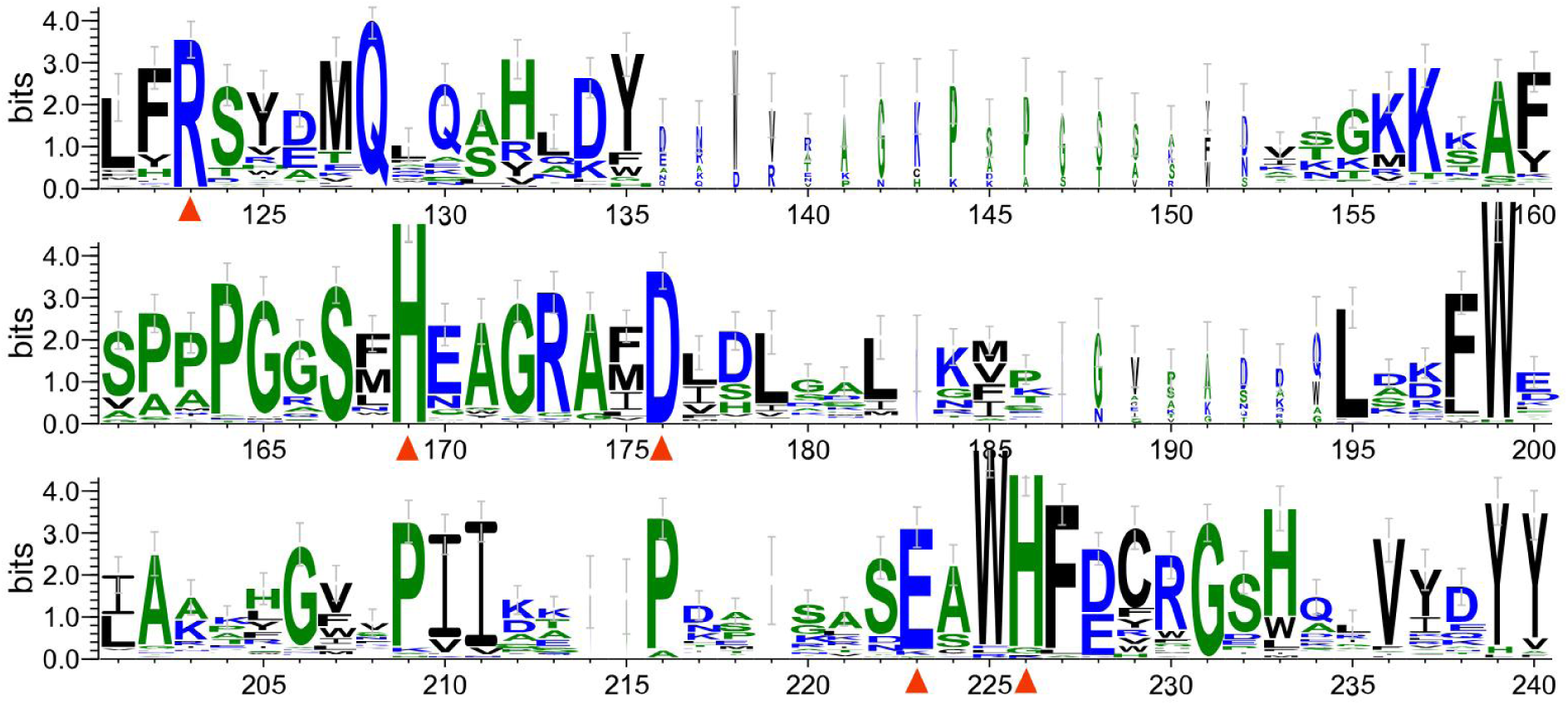
Sequence logos for the multiple sequence alignment of viral L-alanyl-D-glutamate peptidase and homologs (mainly D-alanyl-D-alanine carboxypeptidase) identified from NCBI NR database. The sequence used here are the same as those in the phylogeny shown in Fig. 6. The overall height of the stack represents sequence conservation at each position, while the height of individual symbols within the stack indicates the relative frequency of each amino acid at that position.

